# Presequences of non-imported mitochondrial proteins serve as quality control elements in the cytosol

**DOI:** 10.64898/2026.08.13.744608

**Authors:** Svenja Lenhard, Annika Nutz, Gülsah Göktas, Yury S. Bykov, Markus Räschle, Johannes M. Herrmann

## Abstract

Most mitochondrial proteins are synthesized in the cytosol as precursor proteins with presequences which serve as targeting signals for the mitochondrial matrix, where they are cleaved by the mitochondrial processing peptidase (MPP). In this study, we comprehensively elucidated the role of the presequence and the mature part of mitochondrial precursors in the cytosol, by use of a cytosol-targeted MPP which prematurely processed mitochondrial precursors. Over time, cytoMPP resulted in mitochondrial depletion. However, the cellular response to cytoMPP was surprisingly different to that observed for other models of mitochondrial import inhibition. Cytosolic maturation rendered many proteins stable in the cytosol, indicating that their mature parts lack ubiquitination signals. Accordingly, cytoMPP did not induce the upregulation of the proteasome, which normally is a hallmark of mitochondrial dysfunction. Instead, cytoMPP elicited a heat shock response and impaired the sequestration of precursors in the cytosol. Our observations demonstrate that mitochondrial presequences are more than just address labels. Rather, they play an important role in quality control and orchestrate the cellular response to defects in mitochondrial protein import.

## Introduction

Mitochondria are essential organelles of eukaryotic cells that play central roles in metabolism, energy production and signaling. Mitochondria contain their own genome that codes for a small set of hydrophobic proteins (13 in humans and 8 in baker’s yeast) that are part of the respiratory chain and the ATP synthase complex (Ott et al., 2016). All other proteins (about 1,500 in humans and 900 in baker’s yeast) are nuclear encoded and synthesized in the cytosol as precursor proteins (Morgenstern et al., 2017, Rath et al., 2021). Most of these precursors contain N-terminal matrix targeting signals (MTSs) that serve as address labels to promote their import into mitochondria (Vögtle et al., 2009, von Heijne, 1986). In addition to their role as targeting signals, presequences also serve as binding sites for chaperones and thereby can actively modulate the targeting efficiency of precursor proteins (Rödl et al., 2025, Yan et al., 2026). In specific cases, presequences can even serve as intracellular reporters for import failure: if the precursor of the mitochondrial nucleotide exchange factor Mge1 is not imported it is targeted to the nucleus where its N-terminal segment (i.e. the presequence segment) triggers the mitoCPR stress response (Yuan et al., 2026).

Protein translocases in the outer and inner mitochondrial membranes, also known as TOM and TIM23 complexes, facilitate the translocation of precursors into the matrix (Fig. 1A) (Chacinska et al., 2009, Endo & Wiedemann, 2025). The mitochondrial processing peptidase (MPP) cleaves most precursors to remove the MTS-containing presequence from the mature part of proteins. MPP consists of two essential subunits, Mas1 and Mas2: Mas1 contains the catalytically active zinc-binding site and Mas2 exposes the substrate-binding cavity that is characterized by a glycine-rich loop (Fig. 1B).

**Figure 1:**
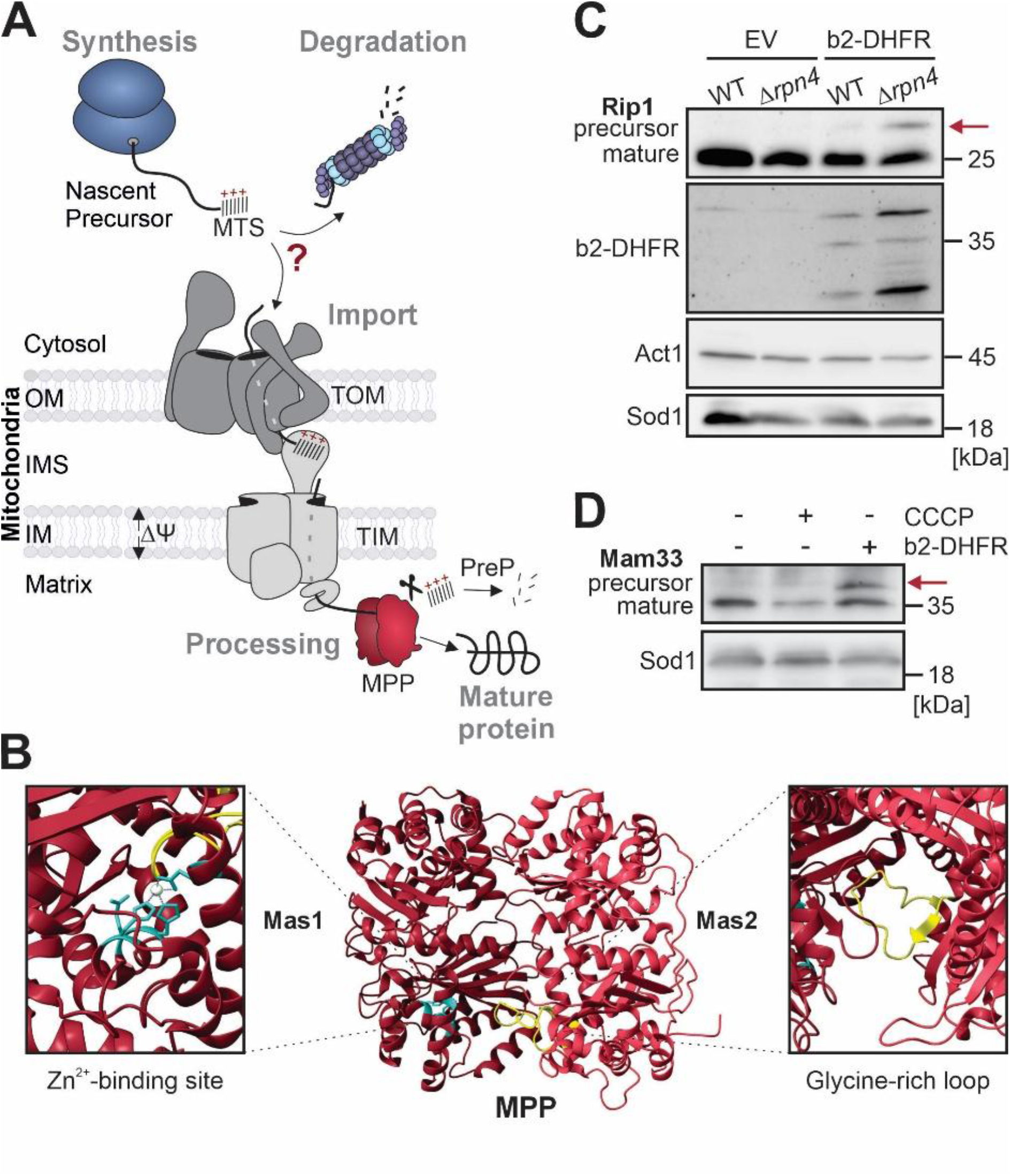
Mitochondrial precursors are proteolytically unstable in the cytosol. (**A**) Schematic representation of the mitochondrial import system. MTS, mitochondrial targeting signal; Δψ, membrane potential; IM, inner membrane; MPP, mitochondrial processing peptidase; OM, outer membrane; PreP, presequence peptidase. (**B**) Structure of the mitochondrial processing peptidase (PDB: 1HR6) (Taylor et al., 2001). The blowouts show the active domains. (**C**) Wild type (WT) and *Δrpn4* cells were transformed with plasmids for the expression of the b_2_-DHFR clogger or with an empty vector (EV). Cells were grown on lactate medium and induced for 6 h with galactose to induce clogger expression. Cells were harvested, lysed and analyzed by Western blotting. The position of the precursor form of the Rieske iron-sulfur protein Rip1 is indicated by an arrow. Act1 and Sod1 are shown as loading controls. (**D**) WT cells with EV (-) or b_2_-DHFR (+) expression plasmids were grown as described for C. Additionally, the cultures were treated with 10 µM carbonyl cyanide m-chlorophenyl hydrazone (CCCP) for 4.5 h after addition of galactose. The position of the precursor form of the matrix protein Mam33 is indicated by an arrow.

If the import of precursor proteins into mitochondria fails, precursors are rapidly degraded by the proteasome (Charmpilas et al., 2025, Rodl & Herrmann, 2023, Song et al., 2021). Therefore, cells respond to the inhibition of mitochondrial import by rapid upregulation of the capacity of the ubiquitin-proteasome system (Boos et al., 2019, Kim et al., 2023, Weidberg & Amon, 2018, Wrobel et al., 2015). Several E3 ligases were shown to cooperate in precursor ubiquitination (Borgert et al., 2022, Samant et al., 2018, Shakya et al., 2021). This redundancy ensures the robust and reliable removal of non-imported mitochondrial proteins. However, the degradation signals (degrons) that are recognized in precursor proteins had not been characterized so far.

Non-imported mitochondrial proteins were proposed to be highly toxic, even though this was only experimentally demonstrated for some non-imported inner membrane proteins, in particular for members of the SLC25 carrier family (Wang & Chen, 2015). These carriers are abundant hydrophobic proteins that are synthesized without presequences and instead use internal targeting signals (Nauerz et al., 2025). Non-imported presequence-containing proteins can be sequestered into transient structures called MitoStores (Amponsah et al., 2025, Bertgen et al., 2024, Ho et al., 2019, Krämer et al., 2023, Samant et al., 2018). Small heat shock proteins induce MitoStore formation, thereby presumably acting as nucleation factors (Peters et al., 2024). Disaggregases, such as Hsp104 in yeast, can remove precursors once stress-conditions calmed down and hand them over to the TOM complex (den Brave et al., 2020, Krämer et al., 2023, Sanchez et al., 1992). Thus, MitoStores serve as dynamic and transient storage particles for mitochondrial precursors in the cytosol.

To better understand the specific roles of the presequences and mature regions of non-imported precursor proteins we generated a cell model in which a cytosol-targeted version of MPP (cytoMPP) was expressed in the cytosol. Expression of cytoMPP removed presequences from mitochondrial precursors. As a consequence, the matured proteins failed to be imported into mitochondria and accumulated in the cytosol. Dynamic isotope labeling allowed us to measure protein stability in these cells. Surprisingly, we found that many mitochondrial proteins remained very stable in the cytosol, indicating that their mature regions lack native degrons. Furthermore, the stress response triggered by these matured proteins strongly differed from that of non-imported precursor proteins. Whereas import failure by cloggers mainly induced the upregulation of the proteasome system, cytoMPP expression triggered a strong heat shock response. Finally, matured proteins were not efficiently sequestered into MitoStores, identifying the presequence as necessary element in MitoStore formation. These observations revealed that presequences carry out complex biological activities: in addition to their role as targeting signals, presequences act as quality control elements which modulate the degradation, folding, and sequestration of proteins. The observations made with our cytoMPP model have strong implications for our understanding of the stress response that is triggered by mitochondrial dysfunction.

## Results

### Mitochondrial precursor proteins are rapidly degraded in the cytosol

The import of proteins into mitochondria can be efficiently inhibited upon depletion of the mitochondrial membrane potential or by expression of clogger proteins that jam the import machinery (Boos et al., 2019, Weidberg & Amon, 2018). For most mitochondrial proteins, only very low amounts of precursors can be detected when the ‘clogger’ cytochrome b_2_(1- 167)-DHFR (or b_2_-DHFR for short), a competitive inhibitor of the mitochondrial import machinery (Boos et al., 2019), is expressed, owing to their rapid degradation by the proteasome (Fig. 1C). Accordingly, *Δrpn4* cells which contain only low proteasome and ubiquitin levels (Xie & Varshavsky, 2001) show increased precursor levels in the presence of b_2_-DHFR.

However, some mitochondrial precursors remain stable in the cytosol and escape degradation; a systematic study using libraries of GFP-tagged proteins estimated that, if protein import into mitochondria was blocked by the use of uncouplers, about 20% of all precursors remained stable in the cytosol (Shakya et al., 2021). The precursor of Mam33 represents one of these stable precursors that can accumulate to larger amounts if protein import is blocked by b_2_- DHFR expression (Fig. 1D). From this we conclude that many mitochondrial precursor proteins contain degrons that allow their efficient removal. These degrons might be part of the presequence and/or the mature segment of the precursor sequence.

### Expression of cyto-MPP produces mature mitochondrial proteins in the cytosol

We next developed a test system to systematically analyze the cytosolic proteolytic stability of the mature regions of mitochondrial proteins (Fig. 2A). To this end, we expressed from a regulatable, galactose-controlled promoter the mature regions of the two subunits of MPP in the cytosol of yeast cells. We therefore cloned the sequences of Mas1 (residue 15 to 463) and Mas2 (residue 10 to 483) into a bidirectional *GAL1/10*-driven expression plasmid fused to Myc or FLAG epitopes, respectively (Fig. 2B). These two proteins, to which we refer as cytoMPP in the following, were not expressed as long as cells were cultured in glucose, raffinose or lactate medium but quickly induced once cells were shifted to galactose (Fig. 2C).

**Figure 2:**
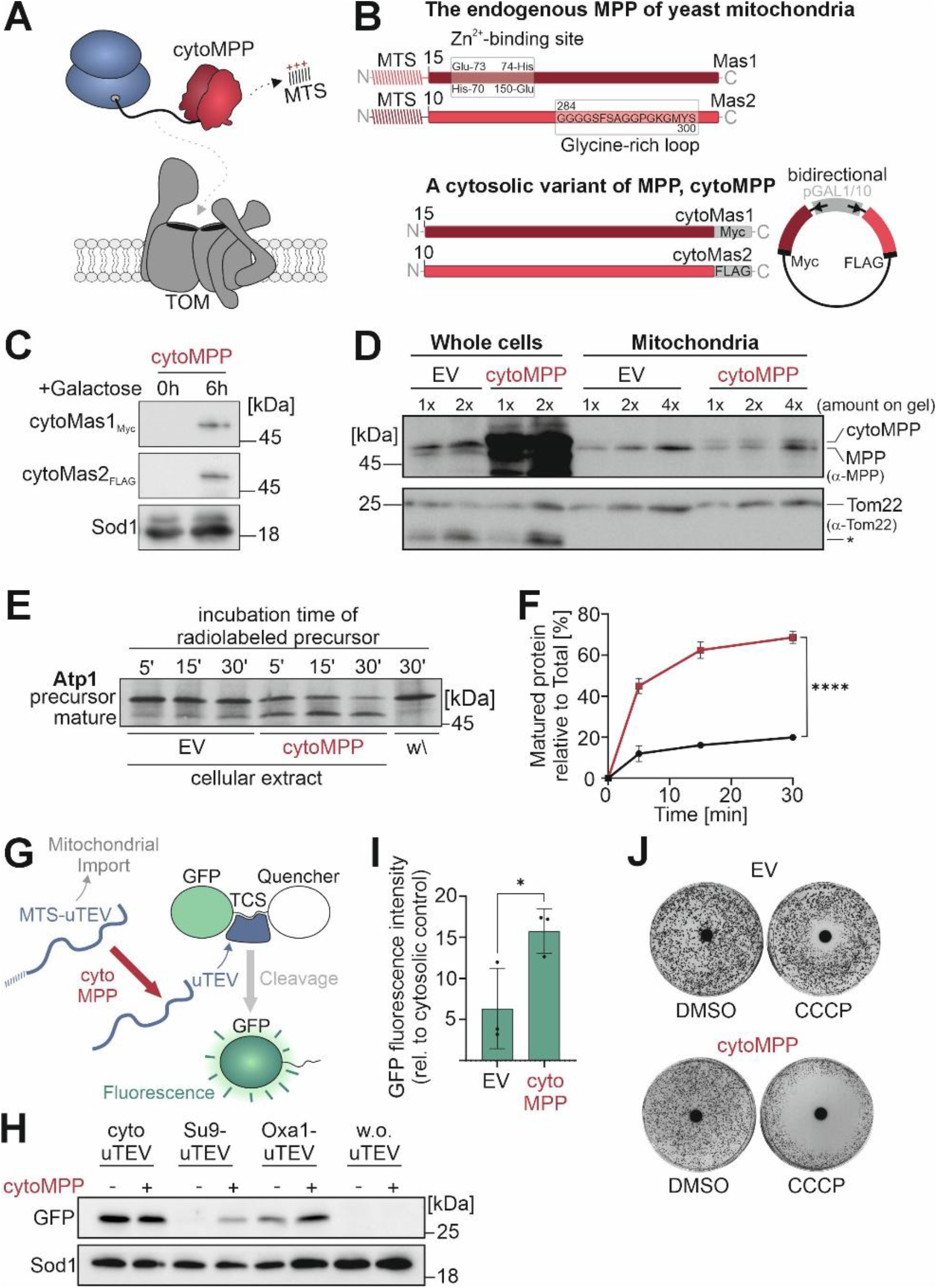
Expression of cytoMPP leads to precursor maturation in the cytosol. (**A**) Schematic representation of the cytoMPP model. (**B**) Overview of the general structure of cytoMPP compared to the endogenous MPP. The proteolytic Zinc-binding site as well as the Glycine-rich loop for substrate recognition are located in the mature parts of Mas1 and Mas2, respectively. The mature forms of the two subunits of MPP, Mas1 and Mas2, are expressed under galactose control from a bidirectional expression plasmid. (**C**) Wild type cells harboring the cytoMPP expression plasmid were grown for 0 or 6 h on galactose-containing medium. Cell extracts were analyzed by Western blotting. Sod1 served as loading control. (**D**) Wild type cells containing the cytoMPP expression vector or an empty vector for control were grown on galactose for 6 h. Cell extracts were directly subjected to SDS-PAGE, or used for the purification of mitochondria that were loaded to the gel. Samples were analyzed by Western blotting. MPP and cytoMPP protein signals were detected using a MPP antibody The signals for Tom22 were shown as control for a mitochondrial protein. (**E**) Radiolabeled Atp1 precursor was incubated with extracts of cytoMPP-expressing cells or control extracts for the times indicated. The control (w\) shows radiolabeled protein that was incubated without cell extracts. Signals were visualized by autoradiography. The positions of precursor and mature forms of Atp1 are indicated. (**F**) The experiment shown in E was carried out three times (n=3) and the signals were quantified. Mean values and standard deviations were plotted. Significance was assessed with a two-way ANOVA statistical test. The P value is indicated as asterisks. **** P≤0.0001 is true for all the time points shown. (**G**) Schematic representation of the *in vivo* assay to monitor cytosolic MPP activity. (u)TEV, Tobacco Etch virus protease; TCS, TEV cleavage site. (**H**, **I**) The indicated proteins were expressed in the cytoMPP or control cells together with the GFP-TCS-Quencher fusion protein. The cleavage of the fusion protein was assessed by Western blotting and detection of the cleaved GFP domain in H, as well as by fluorescence detection from three independent measurements in I. Significance was assessed with an unpaired Student’s t-test. The P value is indicated as asterisks. * P≤0.05 (**J**) Wild type cells harboring cytoMPP expression plasmids or an empty vector were spread on agar plates containing lactate medium with galactose. 10 µl CCCP (10 mM) or DMSO for control was dropped onto a filter disk before plates were incubated for 3 days.

The expression of cytoMPP reached high levels within 6 h of induction, much higher than the endogenous levels of MPP in mitochondria (Fig. 2D). Fractions of isolated mitochondria contained only trace amounts of the cytoMPP bands confirming its cytosolic location (Fig. 2D, lanes labeled ‘Mitochondria’).

Next, we measured the MPP activity of total cellular extracts before and after 6 h of cytoMPP expression (Fig. 2E). We synthesized ^35^S-labeled precursor of the nuclear encoded alpha subunit of the mitochondrial ATP synthase (Atp1), an established substrate of mitochondrial MPP (Vögtle et al., 2009). Upon incubation with cellular extracts, the 53 kDa precursor form of Atp1 was converted into the 47 kDa mature Atp1 protein (Fig. 2E, F). The Atp1 maturation was strongly accelerated upon incubation with cytoMPP-expressing cells, showing that the extracts from these cells contain an about fourfold higher total MPP activity than cells that do not express cytoMPP.

Can the processing activity of cytoMPP also be measured in whole cells? To this end, we designed a reporter strain which expressed a fusion protein consisting of GFP, a tobacco etch virus (TEV) protease cleavage site and a fluorescence quencher domain (Fig. 2G). The quencher prevented GFP maturation so that the strain was non-fluorescent. We further expressed fusion proteins consisting of the MTS of *Neurospora crassa* Su9 or that of Oxa1 fused to an improved variant of the tobacco etch virus protease (uTEV) (Sanchez & Ting, 2020). Since the uTEV protease was rapidly imported into mitochondria, only very low amounts of cleaved GFP (Fig. 2H) and fluorescence (Fig. 2I) were measured in this strain. However, upon expression of cytoMPP, GFP was released and fluorescence was measured. This impressively documented that cytoMPP removed the presequence from the mitochondria-targeted uTEV protease. Upon presequence cleavage in the cytosol, the uTEV domain became active in the cytosol and removed the quencher from the GFP reporter. This fluorescence-based assay was very sensitive and the observed different cleavage efficiencies of constructs with the Su9 and Oxa1 presequences nicely reflected the different import efficiencies of these MTSs (Rödl et al., 2025, Yan et al., 2026).

Interestingly, the expression of cytoMPP made cells hypersensitive to carbonyl cyanide m-chlorophenylhydrazone (CCCP), a compound that retards protein import by uncoupling of the mitochondrial membrane potential (Fig. 2J). This suggests that cytoMPP cleavage can be used as a proxy for import speed or efficiency of individual precursor proteins.

In summary, cytoMPP is a tool to artificially remove presequences from mitochondrial precursor proteins in the cytosol. Thereby it might be used to elucidate the early, cytosolic stages of protein import into mitochondria.

### The expression of cytoMPP leads to mitochondrial protein depletion

Over the last years, many model systems for mitochondrial dysfunction have been described, which despite the common observation of reduced growth and cellular fitness, often considerably differ on a molecular, mechanistic level (Boos et al., 2019, Münch & Harper, 2016, Sutandy et al., 2023, Trifunovic et al., 2004, Wang & Chen, 2015, Wrobel et al., 2015). We therefore studied the effect of cytoMPP in more depth. The induction of cytoMPP expression strongly affected cell growth, particularly on plates (Fig. 3A). High temperature partially alleviated the growth defect. In liquid cultures, cytoMPP expression also retarded cell growth and resulted in reduced cell density maxima, however, over the initial about 12 h of cytoMPP expression, cell growth was rather unaffected (Fig. 3B), suggesting that cytoMPP is not toxic *per se* but causes a problem which manifests over time, presumably by mitochondrial depletion.

**Figure 3:**
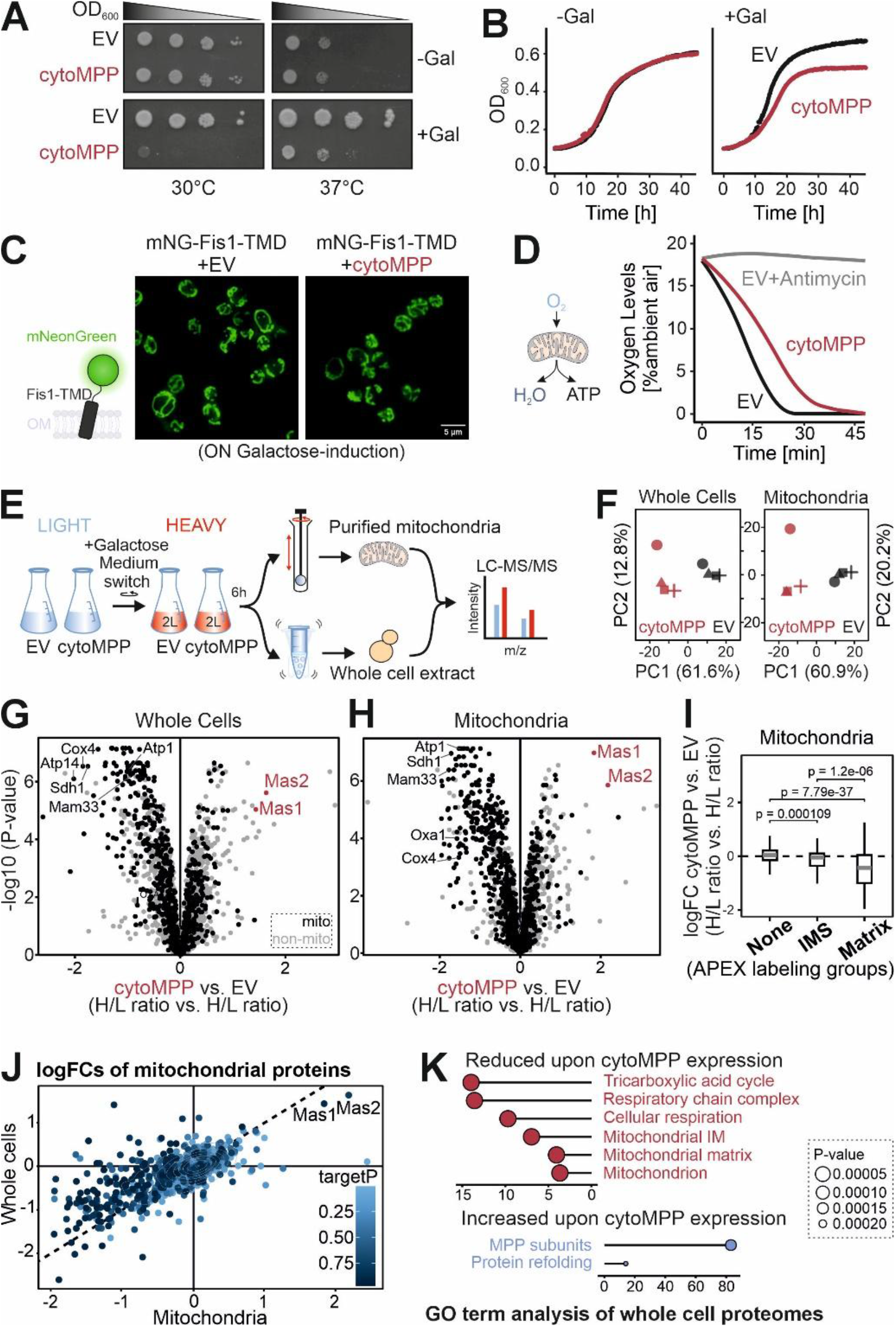
cytoMPP expression leads to the depletion of mitochondrial proteins. (**A**) Wild type cells harboring cytoMPP expression plasmids or an empty vector (EV) were grown to log phase before tenfold dilutions were dropped on lactate (uninduced) or galactose (induced) plates. (**B**) Cell growth of the same strains and media used in A was monitored constantly over 45 h with automated measurements every 10 min using a plate reader. Minimal cultures of 200 µl in a 96-well plate were subjected to constant shaking. Plotted are mean values of three technical replicates. (**C**) A fusion protein of the fluorescent NeonGreen reporter and the outer membrane anchor of Fis1 was expressed in cytoMPP and control cells before mitochondria were visualized by fluorescence microscopy. TMD, transmembrane domain; mNG, monomeric NeonGreen; ON, overnight (**D**) The consumption of oxygen by the indicated cells was monitored. Shown are mean values of four technical replicates. (**E, F**) Schematic representation and principal component analysis of the dynamic SILAC experiment to measure protein levels in cytoMPP and control cells 6 h after shift to galactose medium. (**G, H**) Volcano plots to compare the relative levels of newly synthesized (heavy) proteins in cytoMPP-expressing cells in comparison to the control cells. The positions of some signature proteins are exemplary indicated. (**I**) The levels of many matrix and inner membrane proteins are depleted by cytoMPP expression. The different groups were annotated according to a dataset generated by APEX labeling (Flohr et al., 2025). Significance was assessed using a two-sided Wilcoxon rank-sum test with continuity correction and Benjamini– Hochberg adjustment for multiple testing. (**J**) Correlation plot of protein abundances between whole cell and mitochondrial samples in accordance with the protein specific targetP score. (**K**) GO analysis of protein groups which were particularly enriched or depleted by cytoMPP expression.

To specifically elucidate the consequences of cytoMPP expression, we choose to characterize cells after 18 h of cytoMPP expression. Even though cell growth was already clearly reduced at this time point, the overall morphology of mitochondria remained still unaffected (Fig. 3C). To visualize the mitochondrial network we used a NeonGreen reporter fused to the tail anchor of Fis1, thus a reporter protein that is no substrate of (cyto)MPP.

After cytoMPP expression, cells were also still able to respire efficiently and consumed oxygen at rates similar to those in wild type cells, indicative of a rather unconstrained function of the mitochondrial respiratory chain (Fig. 3D).

We next used a dynamic Stable Isotope Labeling by Amino Acids in Cell Culture (SILAC) strategy to directly assess the cellular proteomes before and after cytoMPP production with high accuracy (Fig. 3E, Table S3). To this end, we grew cells harboring the cytoMPP expression plasmid or an empty vector control in ‘light’ amino acid isotopes in lactate medium (non-inducing conditions) to mid-log phase. Galactose was added to these cultures for 2 h before the cells were reisolated and resuspended in ‘heavy’ [^13^C_6_/^15^N_4_] arginine and [^13^C_6_/^15^N_2_] lysine-containing medium with 0.5 % galactose for continuous cytoMPP induction. The cells were grown for 6 h, during which cell mass roughly doubled so that ‘light’ and ‘heavy’ peptides reached similar levels to ensure accurate quantification. The cells were harvested and either directly lysed or used to purify mitochondria by differential centrifugation. Mitochondrial and whole cell extracts were measured by mass spectrometry. The expression of cytoMPP had a strong consistent impact on the proteomes that were systematically detected in the four biological replicates that we had analyzed for each condition (Fig. 3F, PC1).

From the proteomics samples we calculated the heavy-to-light ratios as this number indicated how much protein was synthesized after the medium shift in comparison to the amount that was present before. Furthermore, we compared the cytoMPP samples to the empty vector control to correct for the effects caused by the carbon source switch. As shown in Figs. 3G and H, we observed a profound depletion of mitochondrial proteins in whole cell extracts as well as in isolated mitochondria.

Among the mitochondrial proteins, matrix and inner membrane proteins showed the strongest reduction, as expected from the fact that the presence of MPP-cleavable presequences is characteristic for these groups (Fig. 3I). Accordingly, the depletion of individual proteins correlated with their targetP score (Emanuelsson et al., 2007), i.e. with the presence of the structural features that are recognized by MPP (Fig. 3J). A GO term analysis revealed many mitochondria-related terms for the group of depleted proteins (Fig. 3K, Table S4). On the contrary, the GO term ‘Protein refolding’ showed an increased abundance.

Thus, in summary, the expression of cytoMPP leads to a depletion of many mitochondrial proteins from cells, consistent with the post-translational import mode of mitochondrial proteins (Eilers & Schatz, 1986).

### The expression of cytoMPP does not trigger the mitoprotein-induced stress response

The accumulation of mitochondrial proteins in the cytosol induces a strong transcriptional response known as mitoprotein-induced stress response or unfolded protein response activated by mistargeting of proteins (UPRam) (Boos et al., 2019, Sutandy et al., 2023, Weidberg & Amon, 2018, Wrobel et al., 2015). In yeast, the expression of clogger proteins which competitively inhibit the mitochondrial import machinery were found to induce three distinguishable response routines on the gene expression level (Fig. 4A): First, the heat shock response was upregulated so that chaperones and other folding factors were increased (Boos et al., 2019, Nowicka et al., 2021). Second, the transcription factor Rpn4 increased the level and capacity of the ubiquitin-proteasome system (Boos et al., 2019, Wrobel et al., 2015). And finally, the targeting of the precursor of the mitochondrial protein Mge1 to the nucleus induces the mitochondrial compromised protein import response (mitoCPR) (Weidberg & Amon, 2018). The transcription factor Pdr3 orchestrates the mitoCPR response.

**Figure 4:**
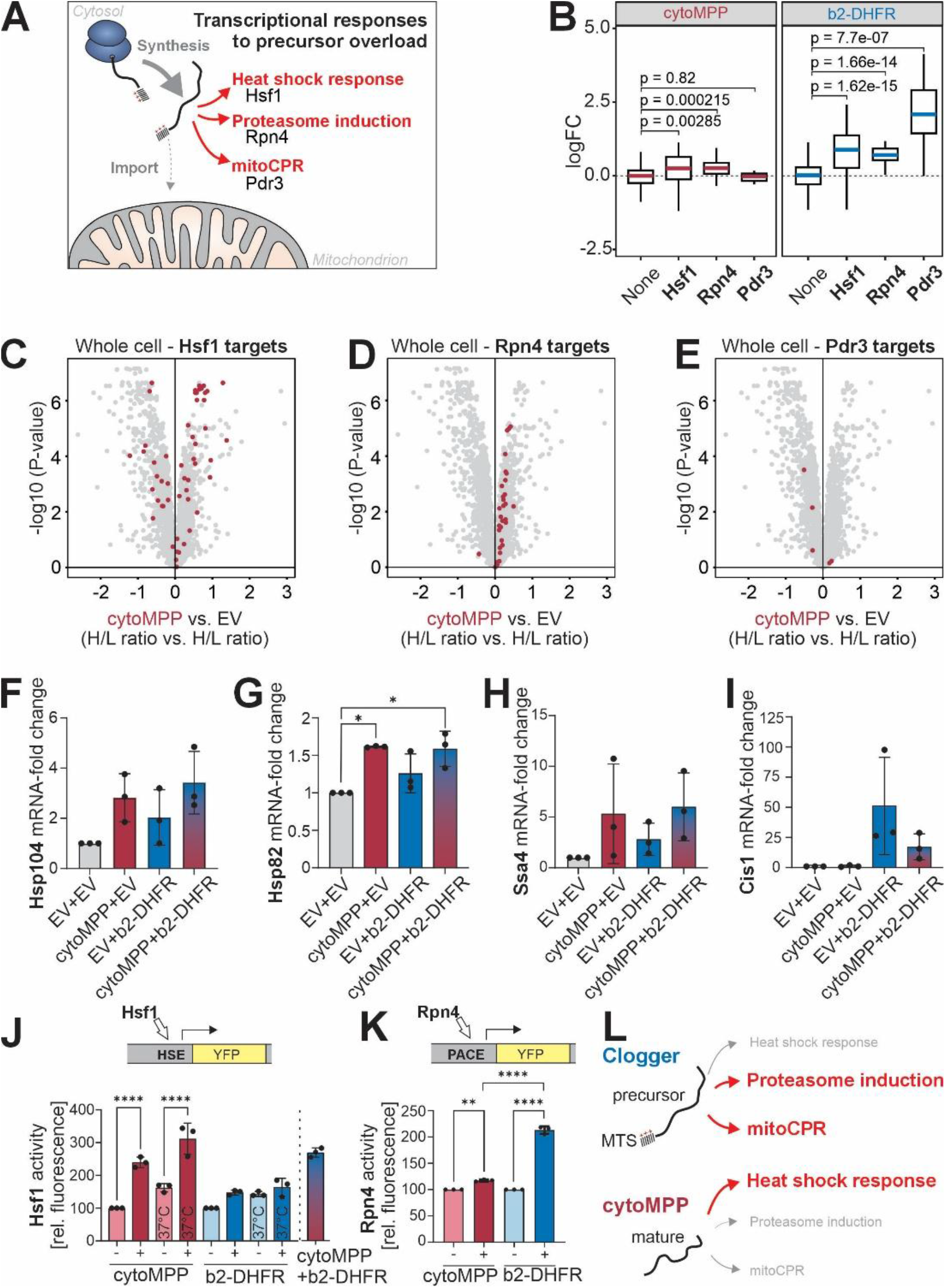
cytoMPP expression triggers a characteristic stress response. (**A**) Schematic representation of the stress response pathways triggered by the aggregation of precursor proteins in the cytosol. (**B**) The relative expressions of the target proteins of the indicated transcription factors are shown. Published data were used for the clogger-induced changes (Boos et al., 2019). (**C-E**) Targets of the transcription factors Hsf1, Rpn4 and Pdr3 were indicated in the volcano plots that show the cytoMPP-induced proteome changes. (**F-I**) Transcript levels of the indicated genes were assessed by qPCR. Shown are mean values and standard deviations from three independent measurements. Significance was assessed using an ordinary one-way ANOVA followed by Tukey’s post hoc test for multiple comparison. The P value is indicated as asterisks. * P≤0.05. (**J, K**) Reporter strains that express yellow fluorescent protein (YFP) under control of a minimal promoter to which a Heat Shock Element (HSE) or a Proteasome-Associated Control Element (PACE) was fused to drive gene expression by Hsf1 or Rpn4, respectively. YFP fluorescence was measured and quantified from three independent repeats. Shown are mean values and standard deviations. Significance was assessed using an ordinary one-way ANOVA followed by Tukey’s post hoc test for multiple comparison. The P value is indicated as asterisks. ** P≤0.01, **** P≤0.0001. (**L**) Schematic representation showing the predominant responses triggered by cytoMPP and clogger expression. See text for details.

When we analyzed the proteomes of cytoMPP-expressing cells we noticed that many chaperones and other proteins that are under control of the heat shock factor Hsf1 were considerably upregulated (Fig. 4C). Apparently, the accumulation of the matured mitochondrial proteins in the cytosol induced a strong heat shock response. In contrast, the levels of proteins under control of Rpn4 and Prd3 were only moderately induced (Fig. 4D, E). Thus, cytoMPP expression clearly shaped the cellular proteome; however, the changes were pronouncedly different to those measured for cells in which the mitochondrial protein import was competitively inhibited by b_2_-DHFR expression (Boos et al., 2019) (Fig. 4B).

To directly elucidate the effects on cellular transcripts, we performed quantitative PCR (qPCR) measurements of cells expressing cytoMPP and b_2_-DHFR individually or simultaneously. First, we measured the levels of mRNAs that are under control of Hsf1 (Fig. 4F-H). *HSP104*, *HSP82* and *SSA4* are well characterized genes that contain heat shock elements as part of their promoters (Yamamoto et al., 2005). All three genes were strongly induced by the expression of cytoMPP; however, b_2_-DHFR expression caused a less pronounced upregulation of the three genes which did not even reach significance levels (Fig. 4F-H). In contrast, b_2_-DHFR expression, but not cytoMPP expression, induced the mitoCPR gene *CIS1* (Fig. 4I). Surprisingly, coexpression of cytoMPP even prevented the b_2_-DHFR - driven *CIS1* induction.

Endogenous genes often have multiple control elements. To exclude indirect effects on such complex endogenous promoters, we made use of a minimalistic reporter system that expresses yellow fluorescent protein (YFP) from a minimal promoter to which either the sequence of the Heat Shock Element (HSE) or the Proteasome-Associated Control Element (PACE) was fused (Boos et al., 2019, Zheng et al., 2016) (Fig. 4J). The induction of cytoMPP induced gene expression with the HSE promoter but had little effect on the PACE reporter (Fig. 4K). On the contrary, b_2_-DHFR expression showed the inverse pattern and mainly induced proteasome upregulation, consistent with previous observations (Boos et al., 2019).

Thus, even though the expression of both b_2_-DHFR and cytoMPP ultimately result in mitochondrial depletion and growth arrest, these two impairments of the mitochondrial import system elicit very distinct transcriptional response programs (Fig. 4L). Apparently, the absence or presence of mitochondrial presequences on accumulating mitochondrial proteins is of considerable physiological relevance: precursor proteins that still contain the MTS strongly induce the expression of the proteasome system and mitoCPR-controlled proteins whereas the accumulation of matured mitochondrial proteins in the cytosol activates the heat shock response, suggesting that the removal of presequences increases the interaction of these matured proteins with cytosolic chaperones (Solis et al., 2018).

### The mitochondrial presequence influences the association with MitoStores

Mitochondrial precursor proteins that accumulate in the cytosol can be sequestered into structures known as MitoStores (also more general CytoQs or Q bodies) (Krämer et al., 2023, Sontag et al., 2017). These foci are bound by the disaggregase Hsp104 and can therefore be efficiently visualized with Hsp104-GFP (Sanchez et al., 1992). To visualize the effect of b_2_- DHFR and cytoMPP expression, we generated strains that expressed Hsp104-GFP constitutively as well as b_2_-DHFR or cytoMPP from a galactose inducible promoter. As shown in Fig. 5A, the induction of cytoMPP induced the formation of Hsp104-GFP foci that were similar to those seen in b_2_-DHFR-expressing cells. Expression of both b_2_-DHFR and cytoMPP also induced Hsp104-GFP foci. For quantification, we identified cells in these images and counted those in which the GFP signals exceeded a Max/Mean threshold of 4.5 (Fig. 5B). The expression of b_2_-DHFR induced MitoStores in about a third of all cells, regardless of whether cytoMPP was co-expressed or not (Fig. 5B). The expression of cytoMPP alone also induced Hsp104-GFP foci and their abundance was similar to the patterns seen in b_2_-DHFR-expressing cells. Just like the MitoStores observed in b_2_-DHFR-expressing cells, the Hsp104-GFP foci induced by cytoMPP were of transient nature as they disappeared once the cytoMPP synthesis was repressed. Within 4 h after cytoMPP expression, the Hsp104-GFP foci became smaller and less frequent (Fig. 5C). Thus, cytoMPP expression induces Hsp104-GFP-positive foci that share the properties described for MitoStores (Krämer et al., 2023).

**Figure 5:**
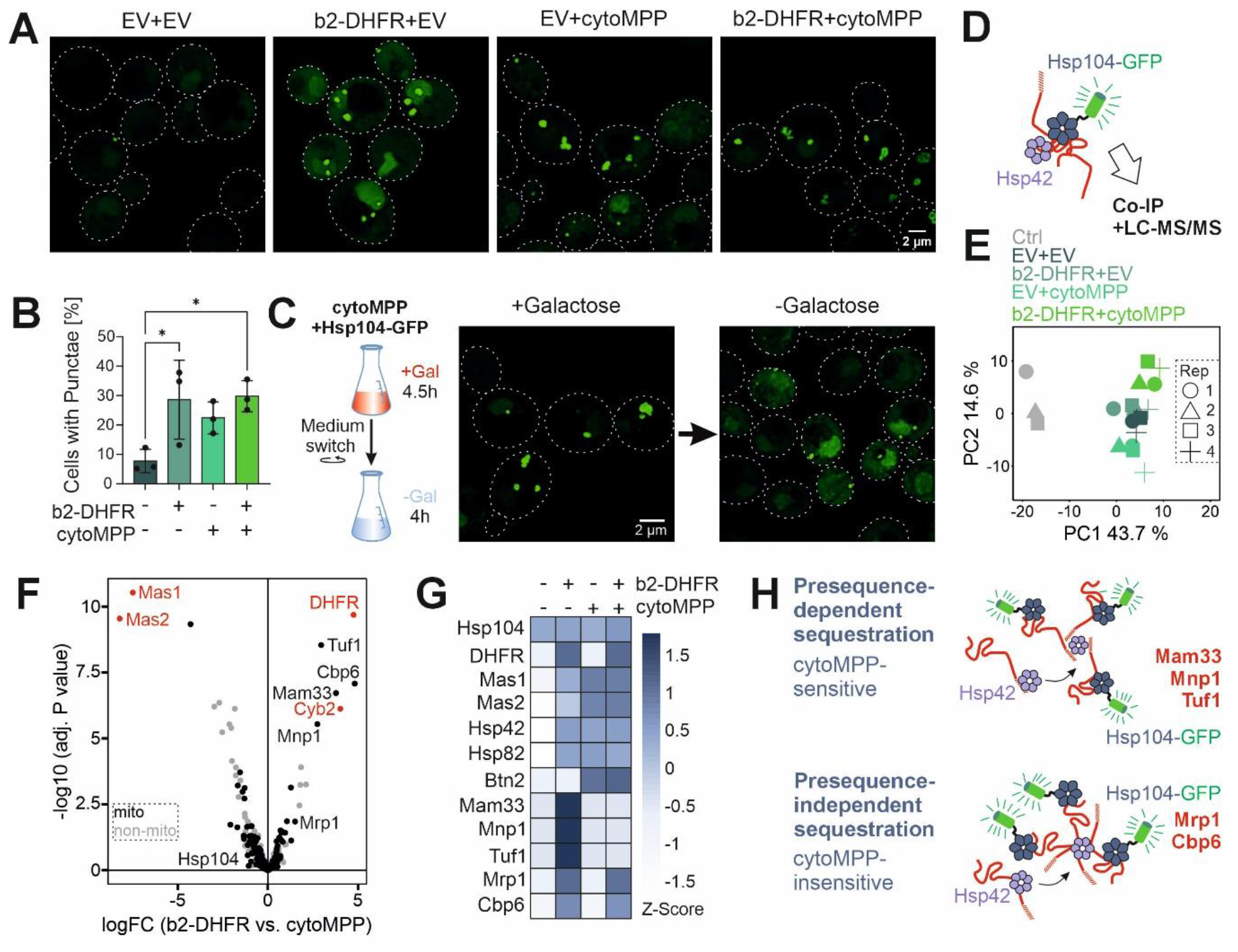
Presequences influence the sequestration into MitoStores. (**A**) Hsp104-GFP was expressed in cells harboring cytoMPP or b_2_-DHFR expression plasmids or empty vector controls. The green foci show Hsp104-containing clusters as previously described for MitoStores. (**B**) The proportion of cells with Hsp104-GFP puncta was quantified. All cells in a bright field image were identified, and the number was set to 100%. The corresponding fluorescent image was screened for cells exhibiting GFP signals exceeding a Max/Mean threshold of 4 and the percentile amount of these cells was calculated for all strains. The bar graphs represent the mean of three independent replicates (n=3) with the respective standard deviation. Significance was assessed by an ordinary one-way ANOVA statistical test. The P value is indicated as asterisks. * P≤0.05. (**C**) Cells were grown in galactose medium for 4.5 h (+ Galactose), reisolated and incubated in galactose-free medium for 4 h (- Galactose) to observe the cytoMPP-dependent formation of Hsp104-GFP foci. (**D**) Schematic illustration of a Co-Immunoprecipitation (Co-IP) of Hsp104-GFP to identify substrates of cytosolic storage granules under the expression of the clogger or cytoMPP. (**E**) Principal component analysis of four biological replicates of whole cell samples subjected to Co-IP followed by mass spectrometry. The different shapes represent each a different replicate. Control (Ctrl) samples without Hsp104-GFP were used to identify proteins that were precipitated nonspecifically. (**F**) Volcano plot of the proteins pulled-down together with Hsp104-GFP under the expression of b_2_-DHFR vs. cytoMPP. The positions of some signature proteins are exemplary indicated. (**G**) Heat map of proteins identified in E and F. Hsp104- GFP substrates can be divided into three groups depending on their presence in Hsp104-GFP foci under the condition of clogger, cytoMPP or simultaneous expression. (**H**) Schematic representation of MitoStore formation dependent or independent of the presequence. The recruitment of proteins by Hsp42 appears to depend on their binding to the presequence or the mature part of precursor proteins.

Next, we identified the Hsp104-GFP-bound proteins in these cells by label free mass spectrometry. We used the same protocol that was used before for the characterization of MitoStores that were formed in the presence of b_2_-DHFR (Krämer et al., 2023).

To this end, we expressed cytoMPP in the absence or presence of b_2_-DHFR in Hsp104-GFP-containing cells. Wild type cells served as control (Fig. 5E, Ctrl). After 6 h of galactose-induction, Hsp104-GFP was purified on magnetic nanotrap beads and the bound proteins were analyzed by mass spectrometry. The principle component analysis shows that the presence of Hsp104-GFP caused a consistent profile as expected (Fig. 5E). Surprisingly, the proteins that were induced to bind to Hsp104-GFP by b_2_-DHFR and cytoMPP considerably differed (Fig. 5F, G, Table S5). Whereas chaperones such as Hsp42 and Hsp82 were found upon both conditions, suggesting that both types of mitochondrial import impairment induce Hsp42- promoted aggregate binding, several MitoStore clients were only found as long as cytoMPP was not expressed. These proteins included Mam33, Mnp1 and Tuf1 for which the presequence seems to be a prerequisite for MitoStore incorporation (Fig. 5H). Other proteins, such as Mrp1 and Cbp6 were found to be associated with Hsp104 upon b_2_-DHFR expression regardless of whether cytoMPP was present or not. In summary, we found that the removal of presequences from precursors can induce the transient formation of Hsp104-bound foci. However, their constituents differ to some degree from the Hsp104-bound proteins that are found after expression of the clogger, showing that the presence or absence of presequences influences the association of these proteins with cytosolic protein aggregates.

### The removal of presequences can protect against proteasomal degradation

The mitochondrial protein import system is under proteasomal control. If protein import fails, most mitochondrial precursors are rapidly degraded in the cytosol (Shakya et al., 2021). However, it is unclear whether the degradation signals of precursor proteins reside in the presequences, the mature parts or in both. Our cytoMPP model for the first time allows it to assess the proteolytic stability of mature regions of mitochondrial proteins systematically. The removal of presequences by cytoMPP leads to the accumulation of matured proteins in the cytosol where they are under proteolytic control of the ubiquitin-proteasome system (Fig. 6A). Stable proteins are expected to accumulate in the cytosol, outside of mitochondria, whereas rapidly degraded proteins are expected to be depleted efficiently. Based on this assumption, we calculate a depletion score which ranked mitochondrial proteins on basis of their cytoMPP-induced accumulation outside of mitochondria (Fig. 6A (right), B, Table S7). This revealed that many of the cytoMPP-cleaved proteins are stable in the cytosol. The mitochondrial termination factor Mtf2, the Lon-type protease Pim1 or the AAA chaperone Hsp78 are examples of this group (Fig. 6C, Table S8). On the other end of the spectrum, proteins such as Nde1 or cytochrome *c_1_* (Cyt1) were found to be strongly diminished by cytoMPP expression. Many of these depleted proteins contained transmembrane domains in consistence with a previous study which found that transmembrane domains can serve as degrons in mitochondrial precursor proteins (Itakura et al., 2016) (Table S7). One might expect that the probability of the presence of degrons correlates with the lengths of proteins if they were stochastically distributed. This was, however, clearly not the case (Fig. 6D). Rather, many long and multidomain proteins accumulated in the cytosol after cytoMPP cleavage. Potentially, the absence of degrons from the mature regions might prevent the unproductive ubiquitination of translocation intermediates, explaining why evolution might have counterselected against such destabilizing elements in mitochondrial precursors.

**Figure 6:**
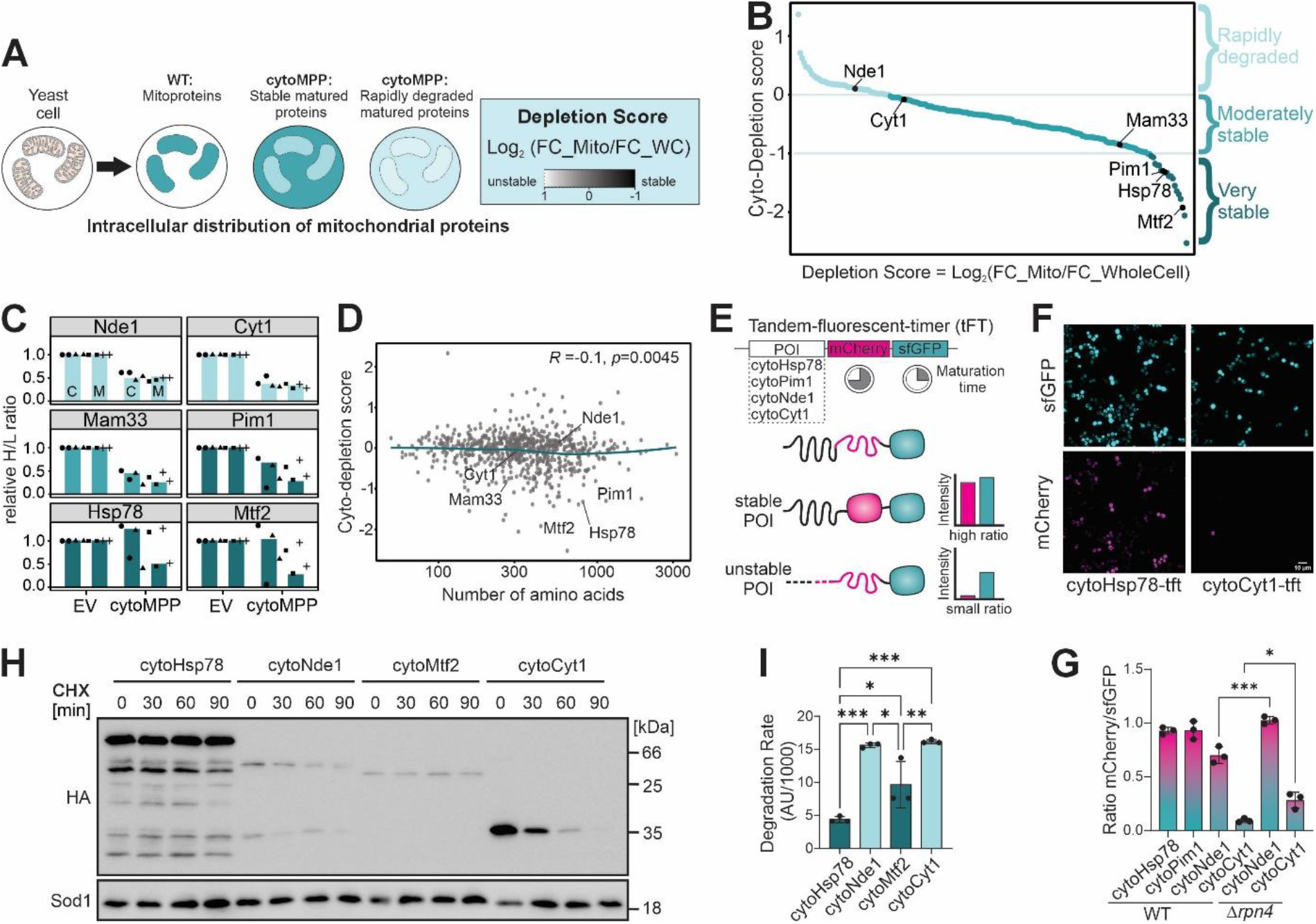
cytoMPP expression allows to measure protein stability of mature mitochondrial protein sequences. (**A**) Schematic protein distribution in the different mutants analyzed. The intracellular accumulation of a presequence-containing mitochondrial protein is schematically depicted in different shades of blue, where lighter shades indicate low abundance. The depletion score can be calculated on basis of the intracellular protein distribution. (**B**) Depletion scores of mitochondrial proteins. Examples are highlighted. The full dataset is provided in Table S7. (**C**) The relative protein heavy-to-light ratios are shown for specific examples, normalized to the empty vector controls. See Table S8. (**D**) Depletion scores of mitochondrial proteins were plotted against protein lengths (at log scale). The correlation between depletion score and protein length was assessed using Spearman’s rank correlation coefficient (*R*). (**E**) Schematic depiction of a fluorescence-based timer consisting of a superfolder GFP domain and a slow folding mCherry domain. (**F**) Two examples of timer constructs were imaged via fluorescence microscopy. Please note the difference in mCherry signal intensities between the two proteins fused to the tandem fluorescent timer. (**G**) Fluorescent signals of the indicated strains were measured with a fluorescent plate reader. Shown are mean values and standard deviations for three independent measurements. Significance was assessed using an ordinary one-way ANOVA test followed by Tukey’s post hoc test for multiple comparison. The P value is indicated as asterisks. * P≤0.05, *** P≤0.005. (**H**) Wild type cells expressing the mature regions of the indicated proteins fused to an HA epitope were expressed from a constitutive promoter. Protein synthesis was blocked by addition of 100 µg/ml cycloheximide. After different times, cells were harvested, lysed and analyzed by Western blotting with the antibodies indicated. Sod1 is shown as loading control. (**I**) Protein stability was assessed by Western blotting as described in G. Shown are mean values and standard deviations of the slope from the degradation up to 60 min of cycloheximide treatment. Three independent experiments (n=3) were quantified. Significance was assessed using an ordinary one-way ANOVA test followed by Tukey’s post hoc test for multiple comparison. The P value is indicated as asterisks. * P≤0.05, ** P≤0.001, *** P≤0.005.

Molecular timers proved to be useful tools to determine the (proteolytic) stability of proteins (Khmelinskii et al., 2012, Kong et al., 2021, Kowalski et al., 2018). These timers consist of two fluorescent proteins in tandem which strongly differ in folding speed (Fig. 6E). In young proteins, only the faster folding protein (in our case superfolder GFP) emits light whereas in older proteins also the slow folding mCherry becomes fluorescent. We expressed the mature parts of several mitochondrial proteins fused to the timer and measured the green and red emission by microscopy (Fig. 6F shown as cyan and magenta, respectively) and in a fluorescence spectrometer (Fig. 6G). The green signal from the superfolder GFP domain was robustly detected in all samples, reaching comparable signals. In contrast, the mCherry signals were much stronger for long-lived mature proteins, such as for Pim1 and Hsp78. Deletion of Rpn4, which diminishes the efficiency of the ubiquitin proteasome system, relatively increased the mCherry signals as expected (Fig. 6G).

Finally, we carried out cycloheximide chase experiments for which we expressed the mature regions of Hsp78, Mtf2, Nde1 and Cyt1 fused to an HA epitope from a constitutive *TPI-* promoter. Translation was stopped by addition of cycloheximide for different times and samples were analyzed by Western blotting (Fig. 6H, I). Hsp78 and Mtf2 remained stable over time, whereas Nde1 and Cyt1 were rapidly degraded. This supports the data of the dynamic SILAC experiment that confirms that the mature regions of many mitochondrial proteins lack degrons.

## Discussion

Most mitochondrial proteins are synthesized with presequences which are necessary and sufficient for mitochondrial targeting (Vögtle et al., 2009, von Heijne, 1986). Presequences share universal structural properties allowing their reliable detection by prediction programs (Emanuelsson et al., 2007). Presequences considerably differ in primary sequence and lengths. Longer presequences often are more efficient and ‘stronger’. These advanced targeting properties might be particularly important to ensure the import of proteins that contain tightly folded structures or hydrophobic transmembrane domains (Rödl et al., 2025, Yan et al., 2026). Such longer sequences can contain binding sites for cytosolic cochaperones of the Hsp90 system and for Tom70 to support the local unfolding of precursor proteins on the TOM pore (Backes et al., 2021, Faou & Hoogenraad, 2012, Rödl et al., 2025, Stan et al., 2000, Yamamoto et al., 2009, Young et al., 2003).

Our observations with the cytoMPP model system suggest that presequences have important functions beyond protein targeting. The expression of cytoMPP caused mitochondrial depletion, similar to what had been observed for the expression of clogger proteins (Boos et al., 2019, Weidberg & Amon, 2018) or for mutants of the mitochondrial import system (Oeljeklaus et al., 2024, Wrobel et al., 2015). However, the cellular response to cytoMPP was surprisingly different to previously used models of mitochondrial dysfunction. Apparently, it is of major physiological consequence whether mitochondrial proteins accumulate in the cytosol with or without their presequences, pointing to a major role of presequences for the stress response of cells (Fig. 7). These observations indicate important functions of mitochondrial presequences beyond their role in protein targeting.

**Figure 7:**
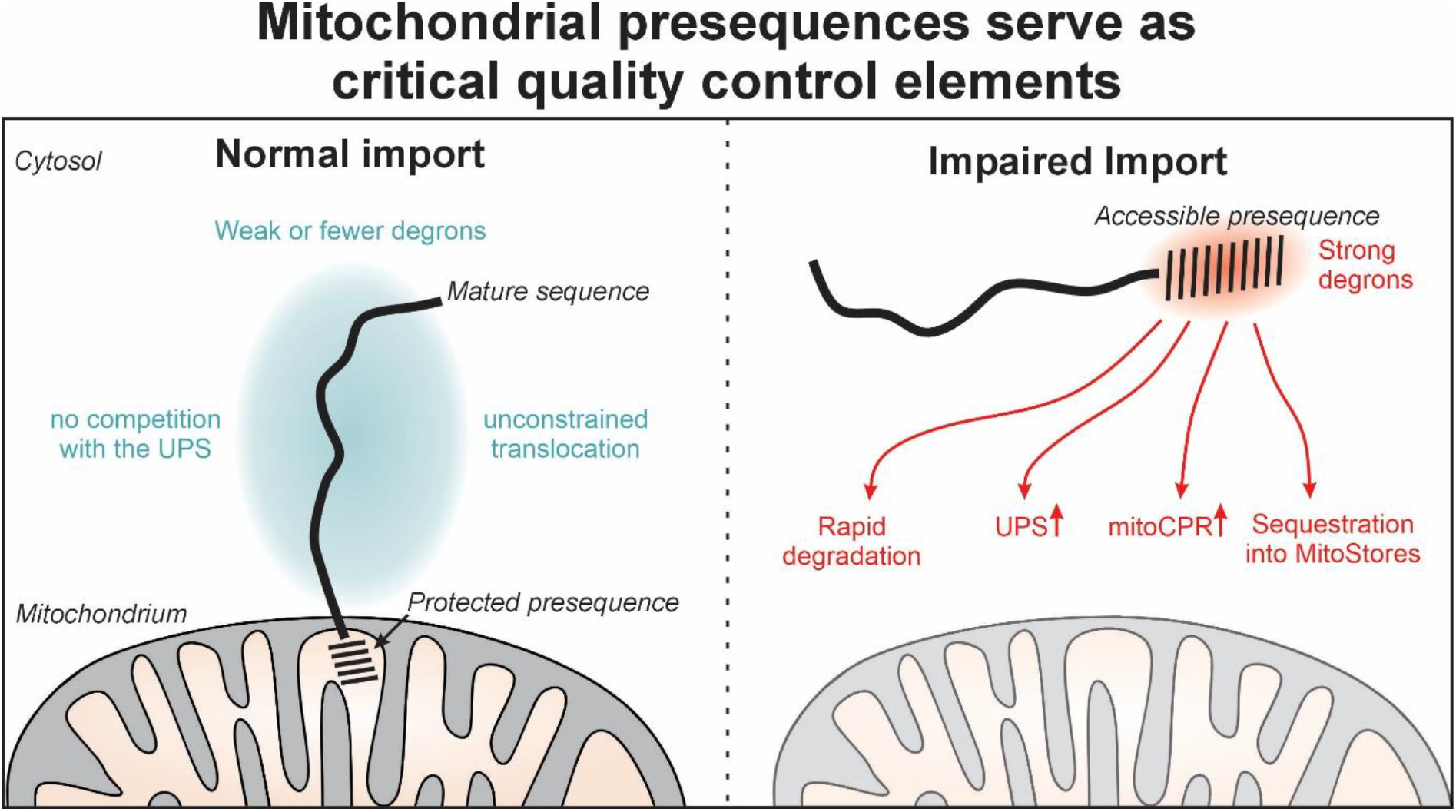
Presequences serve as critical quality control elements. Schematic overview about the presequence-dependent responses described in this study. See text for details.

First, the removal of presequences from mitochondrial precursors caused the stable accumulation of many of these proteins in the cytosol. This shows that many mitochondrial proteins lack degrons in their mature part. The rapid depletion of non-imported precursors seems therefore to be driven by presequences. Interestingly, a large ubiquitination complex was recently discovered in the cytosol of human cells, called the silencing factor of the integrated stress response (SIFI) or UBR4 complex (Grabarczyk et al., 2025, Haakonsen et al., 2024, Yamano & Youle, 2013, Yang et al., 2025). SIFI is a central element in the response to mitochondrial import stress which not only facilitates the degradation of mitochondrial precursors, but also that of the HRI kinase that phosphorylates eIF2alpha.. As soon as precursors accumulate in the cytosol upon mitochondrial dysfunction, the HRI kinase is competed away from the SIFI/UBR4 complex which prevents its degradation; the HRI kinase together with its partner protein DELE2 then triggers the mitochondrial branch of the integrated stress response to support mitochondrial recovery (Condon et al., 2021, Fessler et al., 2020, Guo et al., 2020).

How the cytosolic E3 ligases recognize mitochondrial precursor proteins is not well understood. However, the SIFI/UBR4 complex acts via the N-end rule pathway and recognizes mitochondrial presequences (Tasaki et al., 2005, Yamano & Youle, 2013). Studies in yeast also identified a number of ubiquitin ligases that promote the degradation of non-imported mitochondrial precursor proteins, including Doa10, Hrd1, Mdm30, Mfb1, Rsp5, San1, and Ubr1 (Goodrum et al., 2019, Kowalski et al., 2018, Schulte et al., 2023, Shakya et al., 2021, Zoladek et al., 1997). However, the signals that doom mitochondrial precursors to degradation in yeast cells are less clear than in human cells.

Our observation with the cytoMPP model suggests that – in many precursors - the degrons are confined to the presequence. Many proteins with more than one folded domain, such as Hsp78 or Pim1, seem to lack degrons in their mature region. Arguably, the absence of destabilizing sequences prevents ubiquitination of translocation intermediates formed by such proteins that would lead to a clogging of mitochondrial import channels (Kowalski et al., 2018). Based on our results we propose that mitochondrial precursor proteins show a highly asymmetric distribution of stability elements. As a rule, the presequence of mitochondrial precursors serves as degradation signal for several E3 ubiquitin ligases, but in the mature region of many precursors the occurrence of destabilizing degrons is largely reduced.

A second surprise from our cytoMPP model was the observation that these cells did not upregulate the genes for the ubiquitin proteasome system. The induction of the Rpn4-driven response had been the characteristic feature learned from the clogger expression system as well as from temperature-sensitive import mutants (Boos et al., 2019, Wrobel et al., 2015). Rpn4 levels are controlled by a regulatory loop: this transcription factor is efficiently degraded by the proteasome but accumulates once proteasome substrates competitively prevent its breakdown (Xie & Varshavsky, 2001). Since a large fraction of the cytoMPP- matured proteins are spared from proteasomal degradation, they might simply not compete with Rpn4 for proteasomal degradation so that genes with PACE elements remain unaffected.

The third observation we made was that cytoMPP expression did not induce *CIS1* expression which is the signature gene of the mitoCPR response (Weidberg & Amon, 2018, Yuan et al., 2026). Interestingly, cytoMPP expression even suppressed the clogger-driven expression of CIS1, presumably by removal of the presequence from the Mge1 precursor which triggers the mitoCPR response (Yuan et al., 2026).

Finally, we observed that the removal of presequences from precursors influenced their association with MitoStores in the cytosol in line with the idea that presequences support the sequestration of cytosolic proteins (Krämer et al., 2023). This role of presequences in the sequestration of precursors which presumably avoids their nonproductive interaction with chaperones and explains why the cytoMPP expression induced a very strong heat shock response, which was not observed in other models of mitochondrial dysfunction (Boos et al., 2019, Krämer et al., 2023, Wrobel et al., 2015).

Our study impressively shows that presequences are not simply passive segments with address labels. They are sequences of complex function that help cells to orchestrate the complex reactions that cooperate for the surveillance of mitochondrial biogenesis. It will be exciting to analyze the specific relevance of presequences to models of diseases that are caused by mitochondrial dysfunction. Defects in mitochondrial protein import that jeopardize cellular proteostasis are a hallmark of many diseases and of aging (Coyne & Chen, 2026, Kauppila et al., 2017, Marinho et al., 2023). Our observations of an active role of presequences in this context will open a new avenue for research.

## Methods

### Yeast strains and plasmids

The yeast strains used in this study are based on the wild type strain W303 (*MAT a leu2- 3,112 trp1-1 can1-100 ura3-1 ade2-1 his3-11,15*), YPH499 (*MAT a ura3-52 lys2-801 ade2- 101 trp1-Δ63 his3-Δ200 leu2-Δ*) or BY4741 (*MAT a his3Δ1 leu2Δ0 met15Δ0 ura3Δ0*) (Brachmann et al., 1998, Ralser et al., 2012).

To generate a cytosolically localized version of MPP (cytoMPP), the coding regions for Mas1 and Mas2 without the respective presequences and Stop codons (residues 15-463, Mas1; residues 10-483, Mas2) were amplified from purified genomic DNA of W303 cells. The products were ligated into a pESC plasmid before (cytoMas2) and after (cytoMas1) a bidirectional *GAL1/10* promoter via Gibson assembly. EcoRI and SalI restriction sites were used resulting in C-terminal Myc- and FLAG-tagged cytoMas1 and cytoMas2, respectively. To create a genomically integrating mitochondrial fluorescence reporter construct that does not contain a mitochondrial targeting sequence, mNeonGreen was fused to the transmembrane domain of Fis1 via Modular cloning. In a level 1 assembly, equimolar amounts (75 ng) of all parts were used with the NEBridge® Golden Gate Assembly Kit (BsaI-HF® v2, NEB #E1601). The resulting mNeonGreen-Fis1-TMD fusion construct was integrated into the genome of WT cells after digest with NotI using homolog sequences of the 5’ and 3’ ends of the HO locus. For the analysis of cytosolic stability, the coding regions of the mature parts of Hsp78, Pim1, Nde1 and Cyt1 were amplified from purified genomic DNA of BY4742 cells. The products were each cloned into a pMaK3 plasmid in front of the genes encoding for mCherry followed by sfGFP via Gibson assembly. The genes of mature Hsp78, Nde1, Mtf2 and Cyt1 were ligated via classical cloning into a pYX113 plasmid (GAL1/10 promoter) without a Stop codon resulting in HA-tagged proteins.

Yeast strains and plasmids used in this study are described in detail in the Supplemental Tables S1 and S2.

If not otherwise specified, yeast cells were grown at 30°C in yeast full medium containing 3% (w/v) yeast extract peptione (YEP) broth (Formedium LTD) and 2% of the respective carbon source (glucose (D), galactose (Gal), lactate (Lac), ethanol (E), raffinose (Raf)). Strains carrying plasmids were grown in minimal synthetic (S) medium containing 0.67% (w/v) yeast nitrogen base and 2% of the respective carbon source. For plates, 2% of agar was added to the medium.

### Isolation of mitochondria

For the isolation of mitochondria, yeast cells were propagated in selective galactose or lactate media at 30°C to exponential phase. Cells resulting from a 2 l cultivation were harvested (5 min, 2,000 × g, 25°C), washed with ddH_2_O and treated with 2 ml per g wet weight MP1 buffer (10 mM Tris pH unadjusted, 100 mM DTT) for 10 min at 30°C. After washing with 1.2 M sorbitol, cells were resuspended in 6.7 ml per g wet weight MP2 buffer (20 mM potassium phoshpate buffer pH 7.4, 1.2 M sorbitol, 3 mg/g wet weight zymolyase 20T from Seikagaku Biobusiness) and incubated for 1 h at 30°C. Spheroplasts were collected via centrifugation at 4°C and resuspended in 13.4 ml/g wet weight ice-cold homogenization buffer (10 mM Tris/HCl pH 7.4, 1 mM EDTA pH 8, 0.2% fatty acid free bovine serum albumin, 1 mM phenylmethylsulfonyl fluoride, 0.6 M sorbitol). Spheroplasts were disrupted by 10 strokes with a cooled glass potter. Cell debris was removed by several centrifugations at 1,500 × g for 5 min. To collect mitochondria, the supernatant was centrifuged at 12,000 × g for 12 min. Mitochondria were initially resuspended in 2 ml SH buffer (0.6 M sorbitol, 20 mM HEPES/KOH pH 7.4) and, after another centrifugation at 12,000 × g for 12 min, taken up in 200-400 µl SH buffer (volume depends on the size of the pellet). The concentration of the purified mitochondria was adjusted to 10 mg/ml protein. Aliquots were snap-frozen in liquid nitrogen and stored at −80°C.

### Drop dilution assay

To test growth on plates, drop dilution assays were conducted. Yeast cells were grown in synthetic lactate media to mid-logarithmic phase. After harvesting 1 OD_600_ of cells and washing with sterile water, a 1:10 serial dilution (starting OD = 0.5) was prepared. 3 µl of the dilutions were dropped on agar plates containing lactate medium with or without galactose to determine growth differences. Plates were incubated at 30°C and 37°C. Pictures of the plates were taken after 2 to 4 days of incubation.

### Growth curve

For growthcurves, the microplate reader *SPECTROstar^Nano^*by *BMG Labtech* was used. 1 OD_600_ of a yeast culture in exponential phase (OD_600_=0.5-1) was harvested by centrifugation at 16,000 *g* for 5 min. The cell pellet was washed and resuspended in 1 ml sterile ddH_2_O. In a 96-well plate, 180 µl of selective media and 20 µl of cells were mixed to obtain a final volume of 200 µl per well containing a starting OD_600_ of 0.1. Each measurement was performed in technical triplicates. The 96-well plate was covered with an air permeable membrane. The OD_600_ was automatically measured in intervals of 10 min for 72 h at 30°C. Growth curves were plotted in R.

### Halo assay

To test for growth under chemical treatment, so-called halo assays were conducted. Therefore, an OD_600_=1 of a yeast culture in exponential phase was harvested. The cells were washed and diluted in a 1:100 ratio (final OD_600_=0.01). 100 µl of the cell suspension was plated onto selective media plates using glass beads. A sterile filter disc was placed in the center of the plate and treated with the selected chemical. In this study, 10 µl of 10 mM carbonyl cyanide-m-chlorophenyl hydrazone (CCCP) were used, as well as 10 µl of 100% dimethyl sulfoxide (DMSO) for control plates. After 2-3 days of incubation at 30°C, pictures of the area surrounding the filter disc showing no cell growth (”halo”) were taken.

### *In vitro* activity assay

To prepare radiolabeled (^35^S-methionine) proteins for cytoMPP activity experiments, the TNT Quick Coupled Transcription/Translation Kit from Promega (Walldorf, Germany) was used according to the instructions of the manufacturer. To test the enzymatic activity of cytoMPP, the enzyme was isolated together with the cellular proteome in a native glass bead lysis. Therefore, yeast strains were cultured in SLac medium containing 0.5% Galactose at 30°C. Cells were harvested (OD_600_ =5, 4,000 *g*, 8 min), resuspended in 100 µl lysis buffer (250 mM sucrose, 10 mM MOPS/KOH (pH 7.2), 80 mM KCl, 2 mM MgCl_2_, 5 mM KH_2_PO_4_, 0.1% NP40) and kept on ice. To lyse the cells, several glass beads (Ø 1 mm) were added and the mixture was vortexed for 1 min. To separate glass beads and cellular debris from the cytosolic fraction, samples were centrifuged for 30 s at 15,000 *g* followed by 5 min at 30,000 *g* (4°C). The supernatant of the latter centrifugation step represented the cellular extract (kept on ice). The activity assay was started by preincubating 19 µl of each cellular extract or lysis buffer (control) for 2 min at 30°C. To each sample, 1 µl of radiolabeled protein lysate was added and exposed to enzymatic cleavage in the cellular extract. The reaction was stopped after 5, 15 and 30 min by addition of 5 µl 4x reducing Laemmli buffer (2% sodium dodecyl sulfate, 10% glycerol, 50 mM dithiothreitol, 0.02% bromophenol blue, 60 mM Tris/HCl pH 6.8). Samples were heated (3 min, 96°C) and stored at −20°C. After SDS-PAGE and western blot, results were analyzed via autoradiography.

### Fluorescence-based reporter assays

The *PACE-YFP* or the *HSE-YFP* reporter gene was integrated into the *LEU2* locus of the yeast genome. Cells were grown in SLac medium to exponential phase and 0.5% of galactose was supplemented to the cultures for 6 h. 4 OD_600_ of cells were harvested by centrifugation at 6,000 g for 5 min in 15 ml tubes. The cell pellet was resuspended in 400 µl of SLac medium and 100 µl of the cell suspension was distributed in triplicates to flat-bottomed black 96-well imaging plates (BD Falcon, Heidelberg, Germany). To ensure equal distribution of cells on the well-bottom, the imaging plate was centrifuged gently at 30 g for 5 min. The YFP fluorescence was measured using a ClarioStar Fluorescence plate reader (BMG-Labtech, Offenburg, Germany) by setting excitation (497 nm) and emission (540 nm) wavelengths. A WT strain not expressing YFP was used for subtraction of cellular autofluorescence. An EV control of b_2_-DHFR or cytoMPP conditions were used for normalization of PACE-YFP intensities, whereas a 37°C treated control culture was used for normalization of HSE-YFP measured intensities.

For stability measurements of proteins fused to the tandem-fluorescent-timer construct (Khmelinskii et al., 2012), cells were grown in SD medium to exponential phase and 4 OD_600_ of cells were harvested by centrifugation at 6,000 g for 5 min in 15 ml tubes. Preparation of cells in black 96-well imaging plates was done as described before. The mCherry and sfGFP fluorescence intensities were measured using a ClarioStar Fluorescence plate reader (BMG- Labtech, Offenburg, Germany) by setting excitation (570 nm; 470 nm) and emission (620 nm; 515 nm) wavelengths, respectively. A WT strain not expressing the tft construct was used for subtraction of cellular autofluorescence. The ratio of the measured values for mCherry and sfGFP fluorescent intensities was calculated to assess the stability of the fused proteins of interest.

To measure the uTEV cleavage of the GFP-TCS-Quencher protein in presence of cytoMPP, cells were grown in SLac medium to exponential phase and 0.5% of galactose was supplemented to the cultures for 6 h. Preparation of cells in black 96-well imaging plates and the measurement was done as described before and according to the published protocol for the IQ compete assay (Hoffman et al., 2025).

### Cycloheximide chase assay

For cytosolic degradation analysis of different mitochondrial mature sequences, galactose inducible plasmids containing only the mature part of mitochondrial proteins were transformed into WT cells. Cultures in SLac medium were grown until midlogarithmic growth phase and plasmid expression was induced with 0.5% galactose for 4h. 2 OD_600_ of cells were harvested as a control. All cultures were sequentially treated with 100 µg/ml of cycloheximide and 2 OD_600_ of cells were harvested after 30, 60 and 90 min. Whole cell lysates of all samples were prepared and stored at −20°C until analysis via SDS-PAGE.

### Preparation of whole cell lysates

For the preparation of whole cell lysates, 4 OD_600_ of yeast cells in liquid culture were harvested and washed with sterile water. Pellets were resuspended in 100 µl Laemmli buffer (2% sodium dodecyl sulfate, 10% glycerol, 50 mM dithiothreitol, 0.02% bromophenol blue, 60 mM Tris/HCl pH 6.8) containing 50 mM DTT and boiled at 96°C for 3 min. Cells were lysed using glass beads of 1 mm in diameter and vortexing for 1 min. Alternatively, cells were transferred to screw-cap tubes containing 1 mm glass beads and lysis was performed using a FastPrep-24 5G homogenizer (MP Biomedicals, Heidelberg, Germany) with 3 cycles of 30 s, speed 8.0 m/s, 120 s breaks, glass beads) at 4°C. 100 µl Laemmli buffer was added and samples were stored at −20°C until analysis via SDS-PAGE.

### Antibodies

Act1, Sod1, Rip1, Su9-DHFR, Mam33, MPP and Tom22 antibodies were raised in rabbits using recombinant purified proteins. The secondary antibodies were obtained from Bio-Rad (Goat Anti-Rabbit IgG (H+L)-HRP Conjugate #172-1019; Goat Anti-Mouse IgG (H+L)-HRP Conjugate #1706516). The horseradish-peroxidase coupled HA antibody was purchased from Roche (Anti-HA-Peroxidase, High Affinity (3F10), #12 013 819 001). FLAG and Myc antibodies were obtained from Sigma Aldrich (#F3165) and Invitrogen (#MA1-16637), respectively. Antibodies were diluted in 5% (w/v) nonfat dry milk in 1x TBS buffer with the following dilutions: Anti-HA 1:1000, Anti-Myc 1:1000, Anti-FLAG 1:1000, Anti-Act1 1:500, Anti-Sod1 1:500, Anti-Su9-DHFR1:500, Anti-MPP 1:500, Anti-Tom22 1:5000, Anti-GFP 1:500, Anti-Mouse 1:10,000 and Anti-Rabbit 1:10,000. The Rip1and Tom22 antibodies were kindly gifted by the labs of Thomas Becker and Chris Meisinger, respectively.

### RNA isolation and real-time quantitative polymerase chain reaction (qRT-PCR)

For total RNA extraction, yeast strains were cultivated in synthetic media to exponential growth phase. 4 OD_600_ of cells were harvested, and the RNA was extracted using the RNeasy Mini Kit (Qiagen) in conjunction with the RNase-Free DNase Set (Qiagen) according to the manufacturer’s instructions. RNA quantification was performed in a two-step RT-qPCR. First, 500 ng of extracted RNA were reverse transcribed into cDNA using the qScript™ cDNA Synthesis Kit (Quantabio). Then, to measure relative mRNA levels, the iTaq Universal SYBR Green Supermix (Bio-Rad) was used with 2 µl of a 1:10 dilution of the cDNA sample. Measurements were performed in technical triplicates with the CFX96 Touch Real-Time PCR Detection System (Bio-Rad). Calculations of the relative mRNA expressions were conducted following the 2-ΔΔCt method (Livak & Schmittgen, 2001). Due to its stability, the housekeeping gene *ACT1* was used for normalization.

### Fluorescence microscopy and image analysis

To compare mitochondrial network morphology of cells expressing cytoMPP to control cells, the mNeonGreen-Fis1-TMD fusion construct was integrated into the HO locus of the genome. Cells were cultured in SRaf medium and were continuously diluted to keep them in exponential growth phase and were induced over-night with 2% galactose. To visualize different fluorescence ratios of proteins fused to a tandem-fluorescent-timer consisting of mCherry and sfGFP, respective yeast strains were transformed with an expression plasmid under the control of the constitutive GAP promoter. Cells were cultured in SD medium and were continuously diluted to keep them in exponential growth phase. For microscopy, cells were transferred in volumes of 30-50 µl, depending on the cell density, into concanavalin A coated wells (384-well glass-bottom plates MGB101-1-2-LG-L from Azenta Lifesciences). After 20 minutes, cells were settled to the bottom of the well and unattached cells were removed by washing twice with 1xPBS. Fluorescence microscopy was performed using the Leica-Thunder imager 3D live cell with the 100x oil objective (Leica Microsystems 0.55 S28 LEICA; Inverse, DMi 8 HC; PL Apo 100x/1.44 Oil CDRR CS, LAS X). Wavelengths of 510 nm (mNeonGreen), 575 nm (mCherry) and 475 nm (sfGFP) were used to excite fluorophores. A DFT51010 filter cube was used with emission filters of 519/25 (mNeonGreen, sfGFP) and 594/32 (mCherry). Images were acquired with exposure times of 150-300 ms and laser intensities of 15-30% in a Z-stack format with a spacing of 0.28 µm. Images were saved as 16-bit images in the lif format with a pixel size of 2048 x 2048 pixels and a physical size of 134.15 µm x 134.15 µm. Afterwards, images were processed using the Leica Applications Suite X program (Ver 3.7.4) and Fiji.

For imaging of Hsp104-GFP foci, cells were grown in selective lactate media and were continuously diluted to keep them in the exponential growth phase. Cultures were supplemented with 0.5% of galactose for 6h to induce b_2_-DHFR or cytoMPP expression. Cells were transferred to concanavalin-A coated wells as described before. Fluorescence microscopy was performed using the Olympus SpinSR10 confocal microscope (model IX83 (Evident Scientific)) equipped with a W1 spinning disk scanning unit (Yokagawa) and with Orca Fusion CMOS camera (Hamamatsu) operated with cellSense Dimension v4.4. The images were acquired with UPLXAPO Oil 60x objective (NA 1.42) with 3.2x magnification changer and SoRa spinning disk. A wavelength of 488 nm was used for GFP fluorophore excitation. Images in transmitted light were also acquired. The emission signal was separated with 420/490/550/650 multiple band dichroic mirror and additionally filtered with a 525/25 emission filter. Images at different Z-levels with 0.28 µm spacing were taken with 70% laser intensity and an exposure time of 200 ms.

Image visualization was performed in Fiji/ImageJ (Schindelin et al., 2012) as follows: A maximum projection of all Z-stack images was done, the background fluorescence was subtracted with the rolling ball set to 50 pixels and an unsharp mask was applied with a radius of 3 and a mask value of 0.6. Brightness and contrast were set to the same settings between samples.

### Quantification of Hsp104-GFP foci

To quantify the Hsp104 foci in each cell the middle sections acquired in transmission light were extracted from Z-stacks and segmented using Cellpose (Stringer et al., 2021). The resulting cell masks and maximum projections of the GFP channel were imported into MatLab R2021b (Mathworks) and the maximum and minimum GFP fluorescence intensity was measured in each cell based on the masks. Then the distribution of mean intensities for all cells was plotted. Based on this, the threshold of 175 intensity units was used to remove cells without GFP expression. We also removed the brightest cells that overexpress Hsp104-GFP based on the 95% percentile. For the rest of the cells the ratio of mean intensity to the maximum intensity was calculated and the distribution of this value plotted on the histogram. Based on the histogram, the threshold value of 4.5 was selected to distinguish cells with foci (ratio > 4.5). The resulting classification was overlayed with the GFP images to verify the correct classification. The number of cells with and without foci was quantified in each sample. The similarity of the data distributions was checked for all the replicates, and the same settings were used for all the replicates.

### Oxygen consumption measurements

To measure mitochondrial fitness of cells expressing cytoMPP, the oxygen consumption rate was measured. The strains were cultured in SLac medium to exponential phase, diluted to OD_600_=0.4 and induced with 0.5% galactose for 6 h. 1 OD_600_ was harvested by centrifugation (13,000 g, 5 min) and was washed with ddH_2_O. The cell pellets were resuspended in 125 µl YP medium supplemented with 2% ethanol (YPE) and a dilution series was prepared to obtain concentrations of 8 OD_600_/ml, 4 OD_600_/ml and 2 OD_600_/ml. 10 µl of each cell suspension was used for the measurement. The Multi-Channel Oxygen Reader “FirePlate-O2” (FP96-O2) by PyroScience GmbH was preheated to 30°C and calibrated to oxygen levels of the ambient air. Oxygen values are reported as % air saturation, where ambient air corresponds to approximately 21% O₂. After addition of the differently concentrated cell suspensions to the wells in quadruplicates, the Rainin MicroPro 300 semi-automated pipettor was used to fill all wells of the oxygen sensor microplate with 250 µl of preheated YPE at the same time. The oxygen consumption measurement was started immediately after and stopped after 45 min. As negative control, additional replicates of wild type cells were treated with 10 µl of 10 mg/ml Antimycin A for 10 min before the measurement to inhibit mitochondrial respiration.

### Dynamic isotope labelling of mitochondrial proteins

For dynamic SILAC mass spectrometry (MS), quadruplicates of YPH499 Δarg4 +pESC EV and YPH499 Δarg4 +pESC-cytoMPP cells were cultured in SLac medium containing light [^14^N_2_, ^12^C_6_]-lysine and [^14^N_4_, ^12^C_6_]-arginine isotopes. Cells were diluted continuously to keep them in the exponential growth phase while increasing the culture volume stepwise up to 2 L. The cultures were treated with 0.5% galactose for 2 h to induce cytoMPP expression, before shifting to medium containing heavy [^15^N_2_, ^13^C_6_]-lysine and [^15^N_4_, ^13^C_6_]-arginine isotopes. For the medium shift, all of the culture volume was harvested in 500 ml bottles by centrifugation (5 min, 5,000 g, RT) and the cell pellet was washed with 40 ml SLac medium without lysine and arginine (5,000 g, 5 min, RT). Cells were resuspended in 100 ml SLac medium containing 0.5% galactose and the previously mentioned heavy isotopes. The OD_600_ of this cell suspension was measured and diluted to an OD_600_=0.5 in a volume of 1.5 L. Cultures were incubated for 6 h at 30°C. For whole cell proteomics, 10 OD_600_ were harvested from each culture by centrifugation (5 min, 5,000 g, RT), washed with 10 ml of ddH_2_O. The cell pellet was resuspended in 1 ml of ddH_2_O, transferred to 2 ml tubes and shock frozen with liquid nitrogen. Samples were stored at −80°C for further analysis. For mitochondria isolation, the residual culture volumes were harvested in bottles as described before, washed with 40 ml ddH_2_O and transferred to 50 ml tubes. Cells were pelleted again to remove the water. The cell pellet was shock frozen with liquid nitrogen and samples were stored at −80°C before performing a standard mitochondria isolation as described before.

### Co-immunoprecipitation of proteins bound to Hsp104-GFP

Co-immunoprecipitation followed by mass spectrometry was used for the identification of substrates of GFP fused Hsp104 upon cytoMPP expression in the cytosol compared to the MitoStore-forming condition of b_2_-DHFR expression (Krämer et al., 2023). W303+pESC EV+ pYX233 EV+ pYX142-Hsp104-GFP as well as the same strain without Hsp104-GFP served as negative and background controls, respectively. The expression of only b_2_-DHFR, only cytoMPP or the combination of both was investigated regarding the substrate spectrum of Hsp104. Therefore, quadruplicates of each strain were cultured in 30 ml SRaf medium and diluted continuously to keep them in the exponential growth phase. Cultures were treated with 0.5% galactose for 4.5 h to induce the expression of b_2_-DHFR and cytoMPP. 20 OD600 were harvested by centrifugation (5 min, 5,000 g, RT), washed twice with ice-cold ddH2O and shock frozen in liquid nitrogen. Samples were stored at −80°C for further analysis. Cell pellets were lysed with 400 µl ice-cold lysis buffer (25 mM Tris/HCl pH7.5, 50 mM KCl, 10 mM MgCl2, 5%(v/v) glycerol, 1% (v/v) Nonidet P-40, 1mM DTT, 1mM PMSF, 1x Complete TM Tablets mini EDTA-free protease inhibitor (Roche), 1x PhosSTOP phosphate inhibitor (Roche)) and 1 mm glass beads in the Disruptor Genie (Scientific Industries) for 10 min at 4°C. Lysates were centrifuged for 5 min at 5000 g and 4°C to remove cellular debris. The supernatant was transferred to a new reaction tube and 600 µl of dilution buffer (10 mM Tris/HCl, 150 mM NaCl, 0.5 mM EDTA, 1x Complete TM Tablets mini EDTA-free protease inhibitor (Roche), 1mM PMSF) was added to neutralize the detergent of the lysis buffer and retain molecular interactions. From here on, all steps were performed at 4°C. Magnetic GFP-trap beads (ChromoTek GFP-Trap® Magnetic Agarose) were washed two times with 500 µl dilution buffer and resuspended again in 550 µl dilution buffer. Beads were distributed to pre-cooled reaction tubes and all of each of the lysed samples was added to the beads. To ensure binding of Hsp104-GFP together with its substrates to the beads, samples were tumbling end-over-end for 1 h. All samples were centrifuged for 2 min at 2500 g and a magnetic tube rack was used to separate the beads from the liquid sample. All of the unbound fraction was removed. The beads were washed three times with 500 µl wash buffer I (150 mM NaCl, 50 mM Tris pH 7.5, 5% (v/v) glycerol, 0.05% (v/v) Nonidet-P40) and 2x with wash buffer II (150 mM NaCl, 50 mM Tris pH 7.5, 5% (v/v) glycerol). For the elution of peptides bound to the beads, samples were incubated with 50 µl elution buffer I (2 M urea, 50 mM Tris pH 7.5, 1 mM DTT, 5 ng/µl Trypsin (Promega, Sequencing Grade Modified Trypsin #V5111) for 1h at RT (1000 rpm) and were vortexed every 20 min. Beads and elution buffer were separated with the magnetic tube rack and the supernatant was transferred to a new reaction tube. The beads were again incubated with elution buffer II (2 M urea, 50 mM Tris pH 7.5, 5 mM chloroacetaldehyde, 5 ng/µl Trypsin) for 30 min at RT (1000 rpm) and were vortexed every 10 min. The supernatant of the second elution step was combined with the first elution and samples were incubated overnight in the dark at 37°C.

### Sample preparation and mass spectrometric (MS) identification of proteins

MS samples were prepared after dynamic isotope labelling according to a published protocol with minor adaptations (Kulak et al., 2014). To analyze the mitochondrial proteome changes upon cytoMPP expression, cell lysates were prepared in 100 µl lysis buffer (6 M guanidinium chloride, 10 mM TCEP-HCl, 40 mM chloroacetamide, 100 mM Tris pH 8.5) using a FastPrep-24 5 G homogenizer (MP Biomedicals) with 3 cycles of 20 s, speed 8.0 m/s, 120 s breaks, glass beads (Ø 0.5 mm) at 4°C. Samples were heated for 10 min at 96°C and afterward centrifuged twice for 5 min at 16,000 g. In between, the supernatant was transferred to fresh Eppendorf tubes to remove all remaining glass beads. Protein concentrations were measured using the Pierce BCA Protein Assay (Thermo Scientific, #23225). For protein digestion, 25 µg of protein was diluted 1:10 with LT-digestion buffer (10% acetonitrile, 25 mM Tris pH 8.8). Trypsin (Sigma-Aldrich #T6567) and Lys-C (Wako #125-05061) were added to the samples (1:50 w/w). Samples were incubated overnight at 37°C and 700 rpm. After 16 h, fresh Trypsin (1:100 w/w) was added for 30 min (37°C, 700 rpm). For mitochondrial samples, a standard mitochondria isolation was performed. The protein concentration of isolated mitochondria was measured using the Bradford Assay (Bio-Rad Protein Assay Dye Reagent Concentrate #5000006). 300 µg of isolated mitochondria was pelleted to get rid of the SH buffer needed for storage at −80°C. The mitochondrial pellet was resuspended in 150 µl lysis buffer (6 M guanidinium chloride, 10 mM TCEP-HCl, 40 mM chloroacetamide, 100 mM Tris pH 8.5) and lysis was performed using the Scientific Industries Disruptor Genie cell disruptor for 5 min at 4°C. Samples were boiled for 10 min at 96°C, centrifuged for 15 min at 30,000 g and 4°C and the supernatant was taken for further MS sample preparation. 25 µg of protein (equals 12.5 µl of the supernatant) was taken for trypsin and LysC digestion equivalent to whole cell samples. All further steps were conducted equally for whole cell and mitochondrial samples. The pH of all samples was adjusted to pH <2 with trifluoroacetic acid (10%) and samples were centrifuged for 3 min at 16,000 g and RT. Desalting/mixed-Phase cleanup was performed with 3 layers of SDB-RPS stage tips (cat 2241). Samples were dried down in speed-vac and resolubilized in 12 µl buffer A++ (buffer A (0.1% formic acid) and buffer A* (2% acetonitrile and 0.1% trifluoracetic acid) in a ratio of 9:1). The following steps of Mass spectrometry measurement were performed by Markus Räschle. 4 µl of the resolubilized peptides were separated on a 50 cm long reverse phase column with an inner diameter of 75 µm packed in-house with C18 material (Dr. Maisch GmbH). Chromatography was performed with an Easy-nLC 1200 System directly coupled to a Q Exactive HF mass spectrometer via a Nanoflex source (Sonation). A non-linear 180- minute gradient of 2-95% buffer b [80% (v/v) acetonitrile, 0.1% (v/v) formic acid] was applied using a constant flow rate of 250 nl/min.

Peptides eluted from GFP-trap beads after co-immunoprecipitation were acidified to a pH<2 using Tris-fluoracetic acid and desalted on in-house-prepared StageTips containing Empore C18 disks (Rappsilber et al., 2007). StageTips were activated with 100 µl methanol and twice with 100 µl buffer A (0.1% formic acid). Acidified peptides were loaded onto the StageTips and washed with 100 µl buffer A. Peptides were eluted by addition of 40 µl buffer B (80% acetonitrile, 0.1% formic acid in MS grad water) and dried using a speed vac. Samples were resolubilized in 9 µl buffer A and 1 µl buffer A* (2% acetonitrile, 0.1% tri-flouracetic acid in MS grad water).

4 µl of the resolubilized peptides were separated on a 50 cm long reverse phase column with an inner diameter of 75 µm packed in-house with C18 material (Dr. Maisch GmbH). Chromatography was performed with an Easy-nLC 1200 System directly coupled to a Q Exactive HF mass spectrometer via a Nanoflex source (Sonation). A non-linear 180-minute gradient of 2-95% buffer b [80% (v/v) acetonitrile, 0.1% (v/v) formic acid] was applied using a constant flow rate of 250 nl/min. MS data were processed using MaxQuant Version 2.6.4.0. (Cox & Mann, 2008). All other parameters for data acquisition and processing can be retrieved from the dataset available at ProteomeExchange (see below).

### Mass spectrometric data analysis

The resulting mass spectrometry data sets were processed using the R programming language (R version 4.5.2; R Core Team (2025). R: A language and environment for statistical computing. R Foundation for Statistical Computing, Vienna, Austria. Available online at https://www.R-project.org) similar to previous studies (Flohr et al., 2025, Rödl et al., 2023). For SILAC and co-immunoprecipitation proteomics, the MaxQuant output data was filtered to remove contaminants, reverse hits and proteins identified by site.

For SILAC proteomics, the samples of isolated mitochondria and whole cell samples were analyzed in parallel using identical parameters: H/L ratio was calculated from the label free quantification (LFQ) values of heavy (H) and light (L) channels and filtered to only include proteins that were identified in at least three replicates (n = 4) which resulted in 2105 (isolated mitochondria) and 2616 (whole cell extracts) robustly identified protein ratios. Ratios were then log_2_-transformed, normalized using median centering for each sample, batch effect was removed using the limma package (Quelle) and missing values were imputed if no ratio was calculated in all replicates of any sample by sampling n = 4 values from a normal distribution (seed = 5846). Mean H/L ratios are listed in Table S3.

For co-immunoprecipitation proteomics, the LFQ values were filtered to only include proteins that were identified in at least three replicates (n 4) of the non-control samples (expressing Hsp104-GFP) which results in 883 robustly identified proteins. Log_2_-transformed LFQ values were normalized using variance stable normalization, batch effect was removed with the limma package (Ritchie et al., 2015) and missing values were imputed if a protein was not measured in any sample by sampling n= 4 values from a normal distribution (seed = 6354413). Exemplary proteins were plotted in a heatmap by averaging the replicates per sample and calculating z-scores (Table S6).

Differential expression analysis was performed for SILAC and co-immunoprecipitation proteomics using limma for the indicated comparison of samples and p-values were adjusted for multiple testing using a Benjamini-Hochberg procedure (Benjamini & Hochberg, 1995). Test results are listed in Table S3 (SILAC) and Table S5 (co-immunoprecipitation). The mitochondrial proteome was labeled using a list derived from published data (Morgenstern et al., 2017, Vogtle et al., 2017). For SILAC results, a depletion score was calculated: Depletion Score = log_2_(fold change mito / fold change WCE).

For principal component analysis, H/L ratios or processed LFQ values were used for singular value decomposition of the SILAC or co-immunuprecipitation data, respectively.

Differential expression results of SILAC proteomics were compared to previously published APEX labeling proteomics (Flohr et al) of wild type cells. For this, proteins were grouped in proteins labeled by IMS-APEX (logFC > 1.8 in the respective dataset) or Matrix-APEX (logFC < −1.8 in the respective dataset) and proteins labeled by neither (“other”). The difference between those groups was determined by a two-sided Wilcoxon rank-sum test with continuity correction and Benjamini–Hochberg adjustment for multiple testing. To compare the SILAC proteomics results of this study with the clogger proteomoics dataset previously published (Boos et al), differential expression analysis results were grouped in Hsf1 target, Rpn4 targets or Pdr3 targets and proteins not associated with either stress response (“none”) in both datasets. The difference between those groups was determined separately for each dataset by a two-sided Wilcoxon rank-sum test with continuity correction and Benjamini– Hochberg adjustment for multiple testing.

Correlation between the mitochondrial and whole cell extract samples of the SILAC proteomics as well as correlation between depletion score and protein length was assessed using Spearman’s rank correlation coefficient, calculated with the stat_cor() function from the ggpubr package.

Gene ontology analysis was performed using the GOrilla tool (Eden et al., 2009) on the whole cell extract samples. All quantified proteins were used as background set and proteins with log_2_ fold change > 0.8 and adjusted p-value < 0.05 (enriched proteins or log_2_ fold change < −0.8 (depleted proteins) were used as target set. Results are listed in Table S4.

## Data availability

All reagents and strains are available at request from the corresponding author (JMH).

The mass spectrometry proteomics data (see also Tables S3 and 5) have been deposited to the ProteomeXchange Consortium via the PRIDE (Perez-Riverol et al., 2019) partner repository with the dataset identifier shown below.

Reviewer account details:

| Proteomics Datasets: | Project accession | Token for reviewer access |
| --- | --- | --- |
| Table S3 | PXD082077 | 72J0ytV9HBbo |
| Table S5 | PXD082293 | ilpZTGyq97nG |

## Author contributions

**Svenja Lenhard**: Conceptualization; Data curation; Formal analysis; Validation; Investigation; Visualization; Methodology; Writing—original draft; Project administration; Writing—review and editing. **Annika Nutz**: Formal analysis; Validation; Investigation; Methodology; Software; Writing—review and editing. **Gülsah Göktas**: Data curation; Validation; Investigation; Writing—review and editing. **Yury Bykov**: Data curation; Validation; Formal analysis; Investigation; Writing—review and editing. **Markus Räschle:** Data curation; Validation; Formal analysis; Writing—review and editing. **Johannes M. Herrmann**: Conceptualization; Formal analysis; Supervision; Funding acquisition; Visualization; Methodology; Writing—original draft; Project administration; Writing— review and editing.

## Acknowledgements

We thank Sabine Knaus and Vera Nehr for technical assistance. This study was financially supported by grants from the Deutsche Forschungsgemeinschaft (HE2803/11-1 and SPP2453 project number 541210481 to JMH), the European Research Council (MitoCyto 101052639 to JMH) and the Landesforschungsiniative Rheinland-Pfalz BioComp.

## Abbreviations

APEX: engineered ascorbate peroxidase-based proximity labeling
cytoMPP: cytosol-targeted mitochondrial processing peptidase
DHFR: dihydrofolate reductase
HSE: heat shock element
GFP: green fluorescent protein
MPP: mitochondrial processing peptidase
MTS: mitochondrial targeting signal
PACE: proteasome-associated control element
SILAC: stable isotope labeling of amino acids in cell culture
uTEV: tobacco virus protease
YFP: yellow fluorescent protein.

